## Supplementary Information for "Connectivity Measures for Signaling Pathway Topologies"

This document contains supplementary figures and tables that are referenced in the main text. The code to generate nearly all figures and statistics is available on GitHub at

`https://github.com/annaritz/pathway-connectivity`.

### List of Tables

S1 Reactome signaling pathways considered for pathway influence analysis . . . 4
S2 SIF binary relations . . . 5

[*]Current Affiliation: Department of Computer Science, University of Maryland College Park, College Park, MD, US

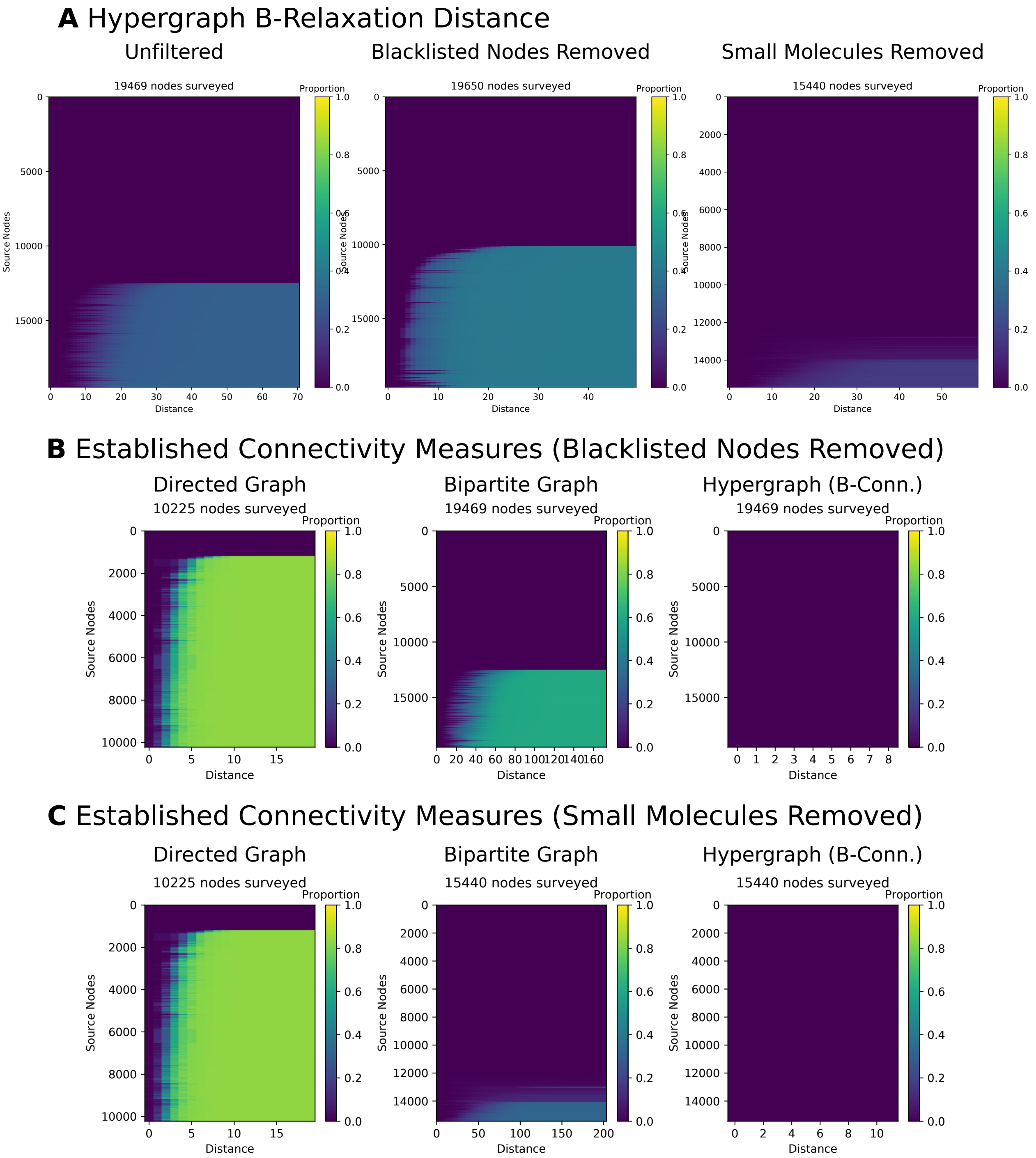

**Fig S1.** Heatmaps showing the effect of filtering pathway representations by blacklisted nodes and small molecules. (A) The proportion of nodes $|B_{\leq k}|$ in the $B_k$-connected set from each source node (rows) for values of $k$ (columns) in the hypergraph. (B) Directed graph connectivity, bipartite graph connectivity, and hypergraph $B$-connectivity for representations with blacklisted nodes removed. (C) Directed graph connectivity, bipartite graph connectivity, and hypergraph $B$-connectivity for representations with small molecules removed.

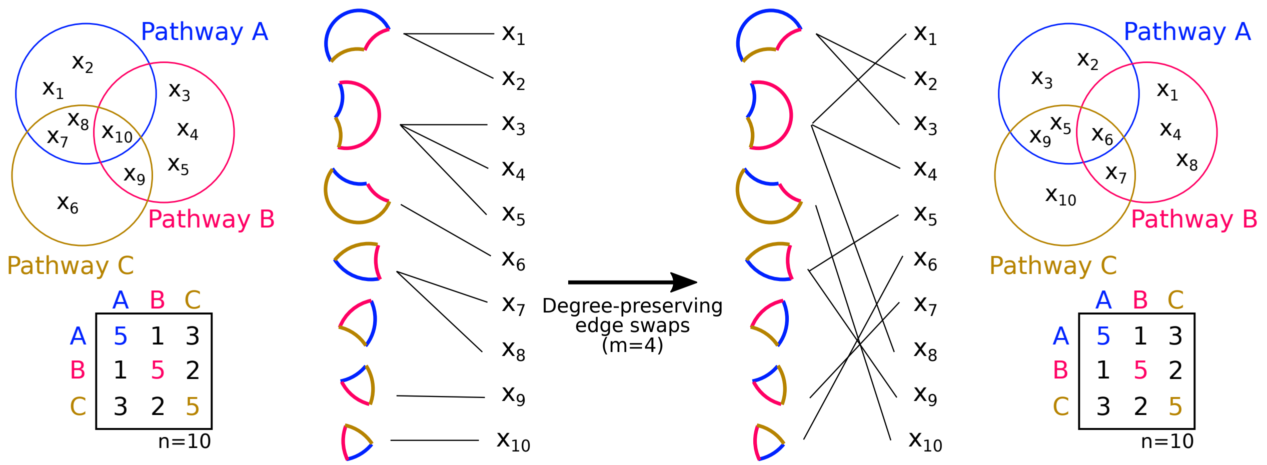

**Fig S2.** Permutation test for pathway membership that preserves initial pathway overlap. The example shows ten molecules ($x_1, \ldots, x_{10}$) that are members of three pathways ($A$, $B$ and $C$). The initial pathway overlap, represented as a Venn diagram, results in the matrix of pairwise overlaps (left). We construct an undirected bipartite graph (we'll call this the *permutation graph* to distinguish this graph from the bipartite graph representation). In the permutation graph, one set of nodes are the molecules and the other set of nodes are all possible overlapping sets except the null set (here, $2^n - 1 = 2^3 - 1 = 7$). Edges in the permutation graph connect molecules to the overlapping set to which they belong. We then perform degree-preserving edge swaps by selecting pairs of edges with different nodes and swapping them (we perform 10,000 swaps in all experiments). We then "piece" the Venn diagram back together, which contains the same number of elements in each overlapping set (resulting in the same pairwise overlaps). With 34 pathways, one would think that the permutation graph is too large; however, we found that there were only 255 non-empty portions of the Venn diagram for the hypergraph/bipartite graph entities (and only 180 for the directed graph entities). The size of the permutation graph also makes the choice of 10,000 swaps for each permutation reasonable.

| Signaling Pathway | Reactome ID | # in Pathway | # in Hypergraph | # in Filtered Hypergraph |
|---|---|---|---|---|
| EGFR | R-HSA-177929 | 164 | 110 | 101 |
| ERBB2 | R-HSA-1227986 | 220 | 118 | 105 |
| ERBB4 | R-HSA-1236394 | 181 | 101 | 92 |
| PI3K/AKT | R-HSA-1257604 | 753 | 357 | 333 |
| MET | R-HSA-6806834 | 240 | 130 | 121 |
| FGFR | R-HSA-190236 | 373 | 232 | 221 |
| ERK1/ERK2 | R-HSA-5684996 | 656 | 310 | 291 |
| IGF1R | R-HSA-2404192 | 160 | 75 | 64 |
| Insulin | R-HSA-74752 | 207 | 90 | 76 |
| Integrin | R-HSA-9006921 | 88 | 66 | 53 |
| GPCR | R-HSA-372790 | 2456 | 1006 | 796 |
| DAG/IP3 | R-HSA-1489509 | 107 | 50 | 32 |
| PDGF | R-HSA-186797 | 244 | 88 | 80 |
| VEGF | R-HSA-194138 | 351 | 215 | 183 |
| NTRKs | R-HSA-166520 | 362 | 205 | 184 |
| Wnt | R-HSA-195721 | 921 | 358 | 326 |
| TNF | R-HSA-75893 | 127 | 97 | 93 |
| PTK6 | R-HSA-8848021 | 187 | 122 | 114 |
| TGFB | R-HSA-170834 | 247 | 179 | 166 |
| TRAIL | R-HSA-75158 | 17 | 13 | 13 |
| FasL/CD95L | R-HSA-75157 | 10 | 10 | 10 |
| Notch | R-HSA-157118 | 505 | 286 | 271 |
| BMP | R-HSA-201451 | 74 | 38 | 36 |
| Activin | R-HSA-1502540 | 44 | 30 | 28 |
| MAPK4/MAPK6 | R-HSA-5687128 | 200 | 113 | 105 |
| NTR | R-HSA-193704 | 245 | 141 | 129 |
| SCF-KIT | R-HSA-1433557 | 122 | 86 | 79 |
| Hedgehog | R-HSA-5358351 | 417 | 190 | 172 |
| Nuclear | R-HSA-9006931 | 431 | 261 | 208 |
| Leptin | R-HSA-2586552 | 54 | 33 | 31 |
| Hippo | R-HSA-2028269 | 79 | 47 | 45 |
| Rho GTPases | R-HSA-194315 | 808 | 330 | 291 |
| MST1 | R-HSA-8852405 | 21 | 12 | 9 |
| mTOR | R-HSA-165159 | 111 | 67 | 57 |

**Table S1. Thirty-four Reactome signaling pathways considered for the pathway influence analysis.** Members that are not part of any hyperedge are ignored from the hypergraph. The filtered hypergraph has removed all small molecules, two forms of Ubiquitinase, and the Nuclear Pore Complex from the hyperedges.

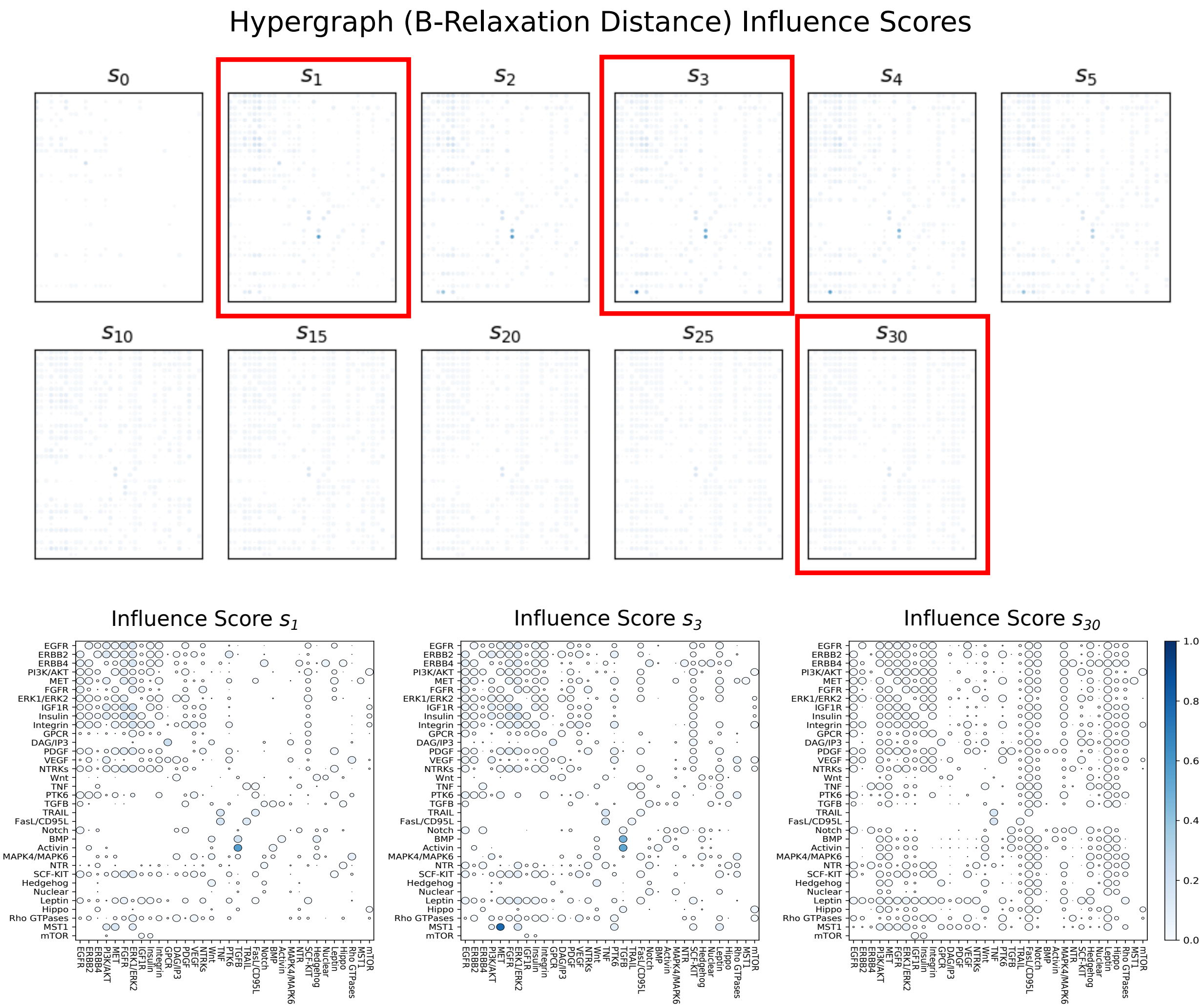

**Fig S3.** Influence scores of pairs of Reactome pathways for the hypergraph at selected values of $B$-relaxation distance $k$. Rows indicate the source pathway $P_S$ and columns indicated the target pathway $P_T$. Color indicates influence score and circle size indicates significance by permutation test (larger circles are more significant). Three selected distances are enlarged (note $s_3$ is also in the main manuscript).

| Binary Relation | # of Relations in Reactome | Conversion Rule |
|---|---|---|
| catalysis-precedes | 263,342 | Directed Edge |
| chemical-affects | 15,473 | Directed Edge |
| consumption-controlled-by | 8,079 | Directed Edge |
| controls-expression-of | 3,730 | Directed Edge |
| controls-phosphorylation-of | 3,184 | Directed Edge |
| controls-production-of | 8,052 | Directed Edge |
| controls-state-change-of | 113,558 | Directed Edge |
| controls-transport-of | 5,232 | Directed Edge |
| controls-transport-of-chemical | 5,105 | Directed Edge |
| in-complex-with | 140,302 | Undirected Edge |
| reacts-with | 1,922 | Undirected Edge |
| used-to-produce | 5,888 | Directed Edge |

**Table S2. Rules for converting SIF binary relations to directed edges.** We ignore the "neighbor-of" binary relation.

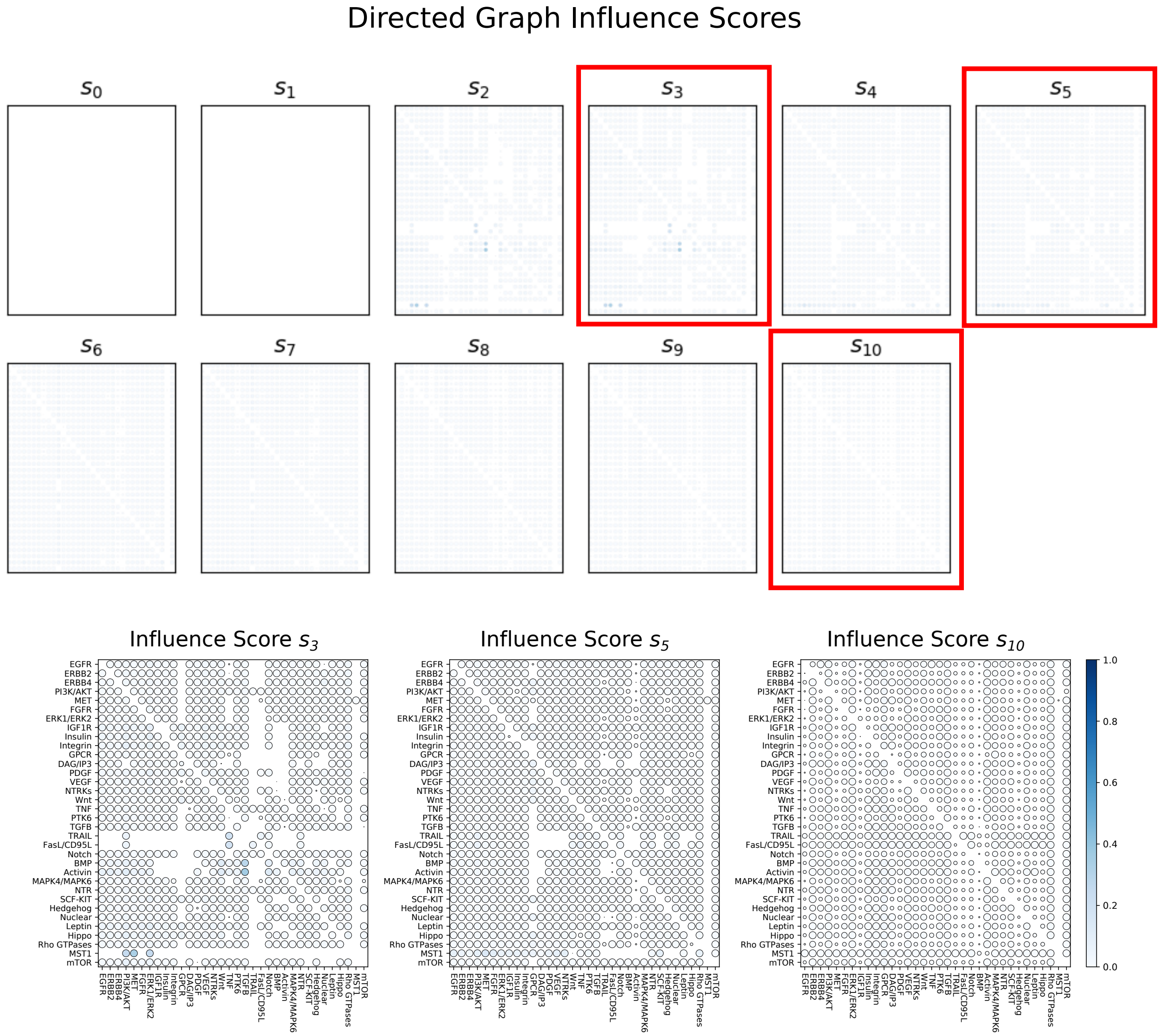

**Fig S4.** Influence scores of pairs of Reactome pathways for the directed graph at selected values of distance $k$. Rows indicate the source pathway $P_S$ and columns indicated the target pathway $P_T$. Color indicates influence score and circle size indicates significance by permutation test (larger circles are more significant). Three selected distances are enlarged. Note that the entities are different for this graph than the hypergraph and bipartite graph, resulting in different initial pathway overlaps.

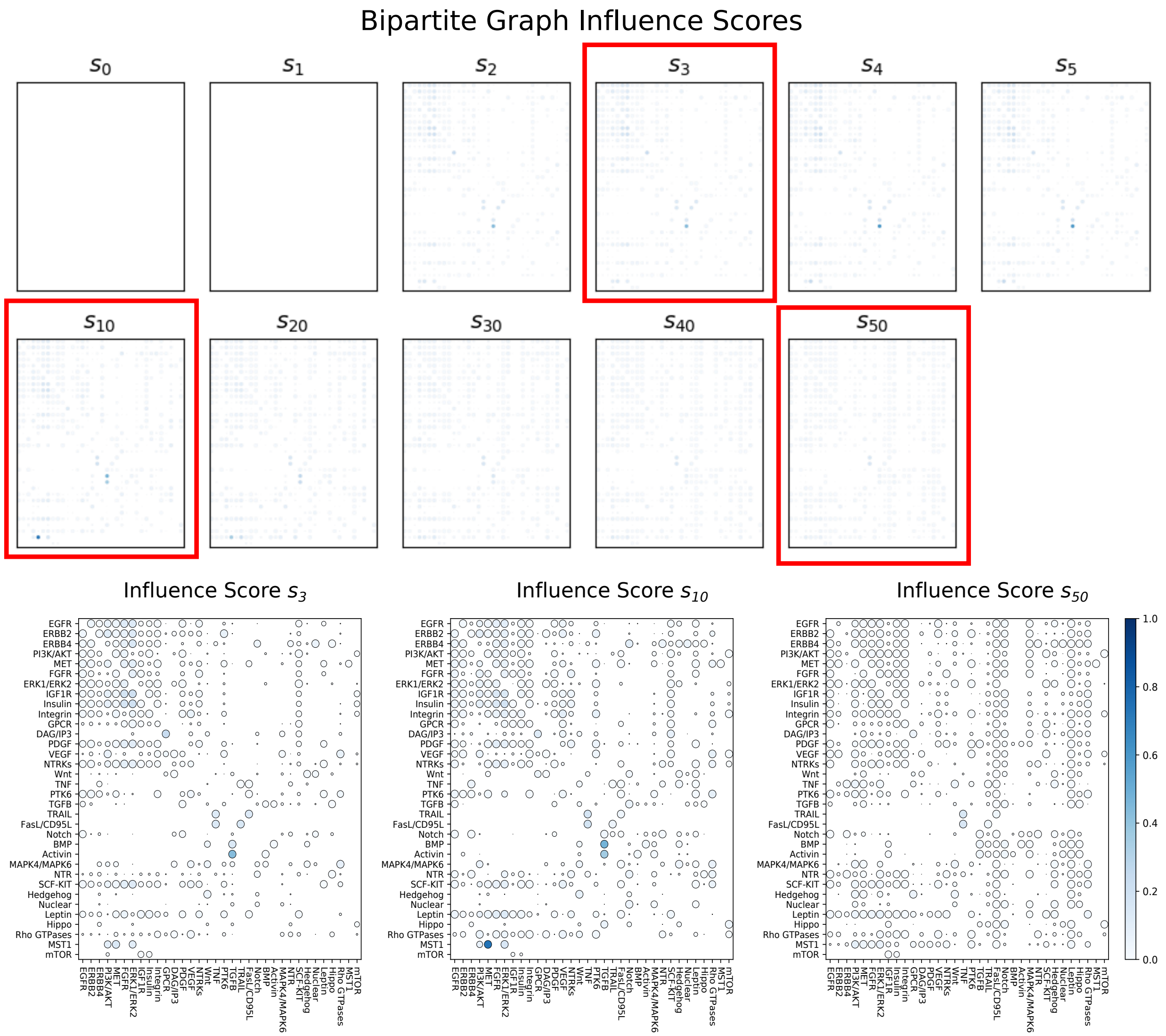

**Fig S5.** Influence scores of pairs of Reactome pathways for the bipartite graph at selected values of distance $k$. Rows indicate the source pathway $P_S$ and columns indicated the target pathway $P_T$. Color indicates influence score and circle size indicates significance by permutation test (larger circles are more significant). Three selected distances are enlarged.

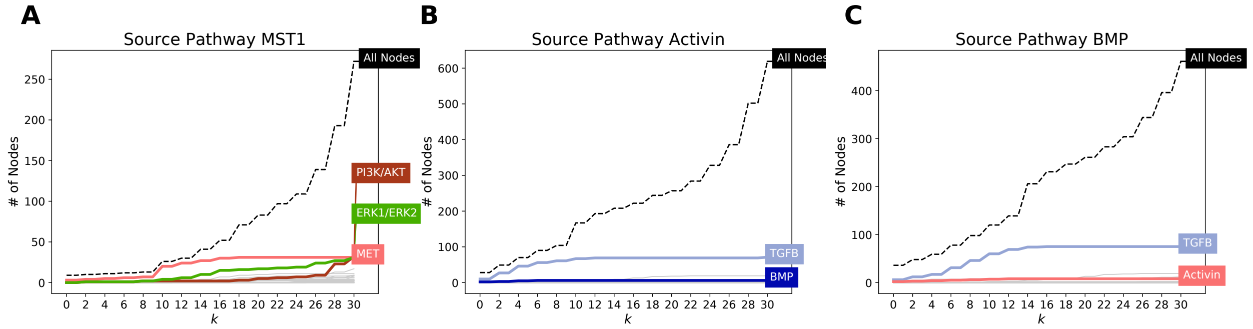

**Fig S6.** textbfA single pathway's influence on downstream pathways in the bipartite graph. Shown are the influence of (A) signaling by Mst1, (B) signaling by BMP, and (C) signaling by Activin. The dashed black line indicates the number of nodes in the source pathway's $B_{\leq k}$ for different values of $k$. There is one line for each of the 33 other target pathways denoting the number of members that appear in $B_{\leq k}$, with selected pathways highlighted in bold.

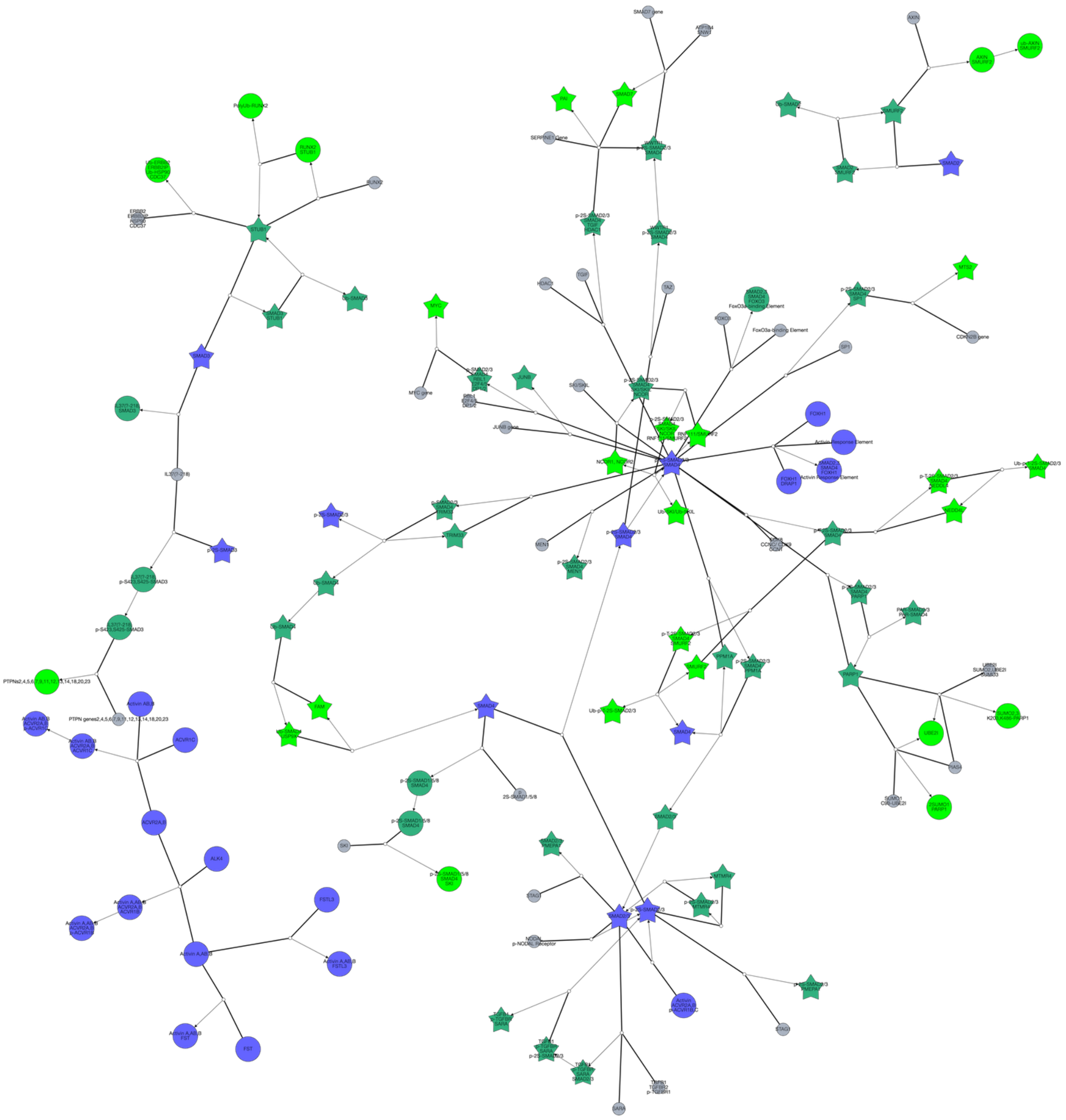

**Fig S7. Hyperedges traversed to compute $B_0, B_1, \ldots, B_4$ from source pathway Activin.** Node colors represent $B$-relaxation distance from $k = 0$ ($B$-connected set, blue) to $k = 3$ (bright green). Gray nodes are entities that are not in the $B_k$-connected set but are involved in traversed hyperedges. Star-shaped nodes are members of the TGF$\beta$ pathway. This network is available on GraphSpace at `http://graphspace.org/graphs/26756?user_layout=6713`.

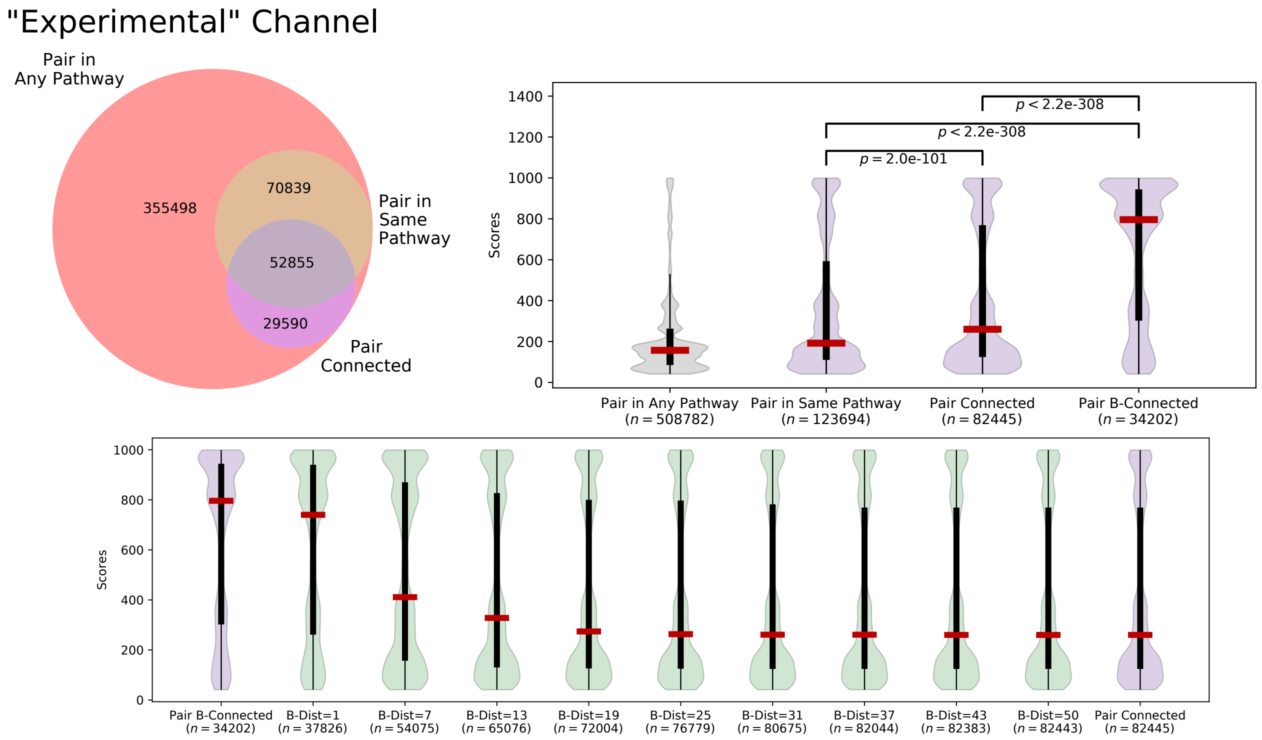

**Fig S8.** STRING interactions within Reactome for "experimental" interactions. In addition to the components of Fig. 7, the bottom violin plot shows the distributions of interaction scores for selected $B$-relaxation distance thresholds.

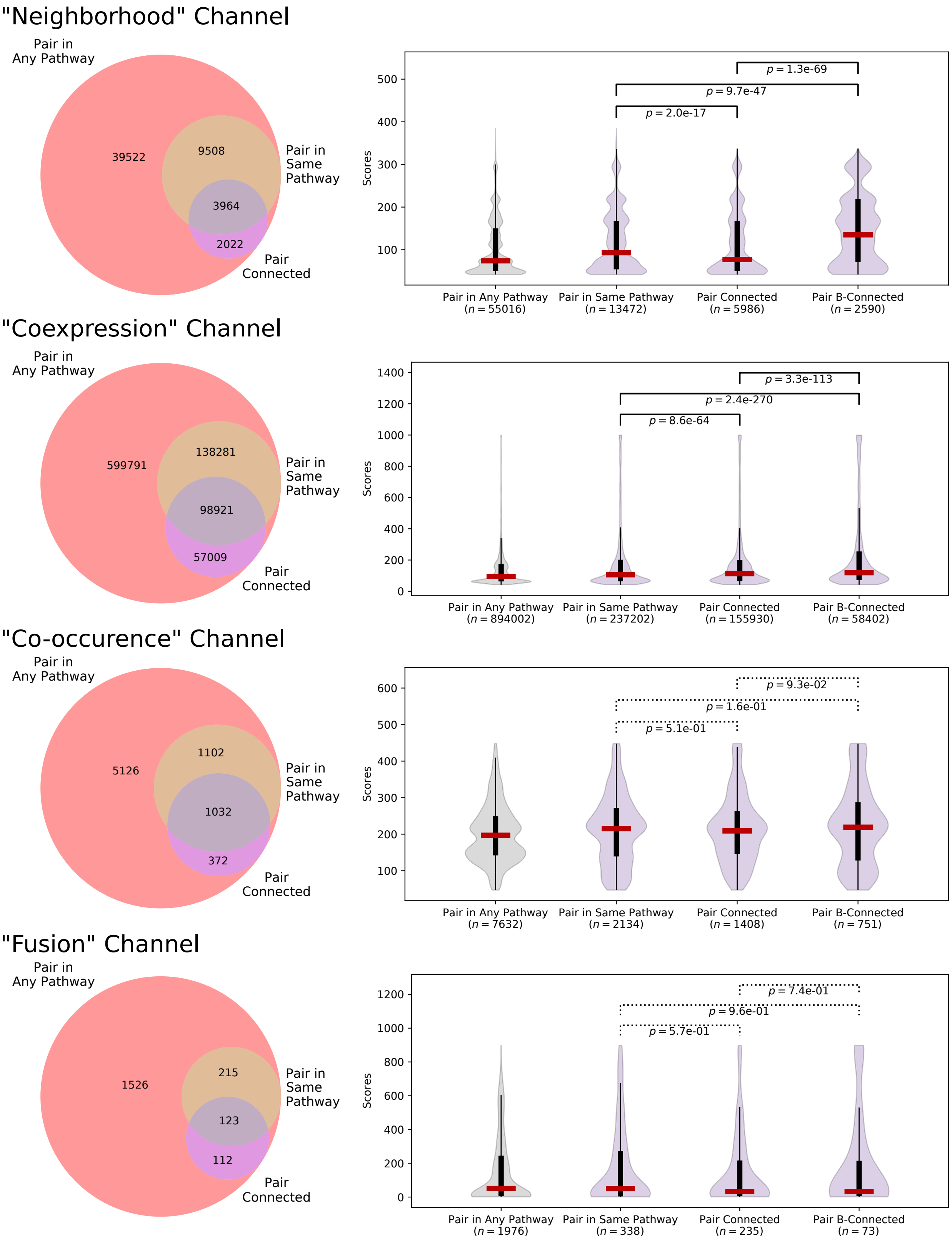

**Fig S9.** STRING interactions within Reactome for the remaining evidence channels not shown in Fig. S8 or Fig. 7. The Venn diagram shows the overlap of interactions where the nodes appear in any Reactome pathway, appear in the same Reactome pathway, or are connected in the bipartite graph. The violin plot shows the distributions of interaction scores (which range from 1 to 1000) for different sets of interactions (median and percentiles shown; Kruskal-Wallis $p$-values less than 0.01 are shown with a solid line).

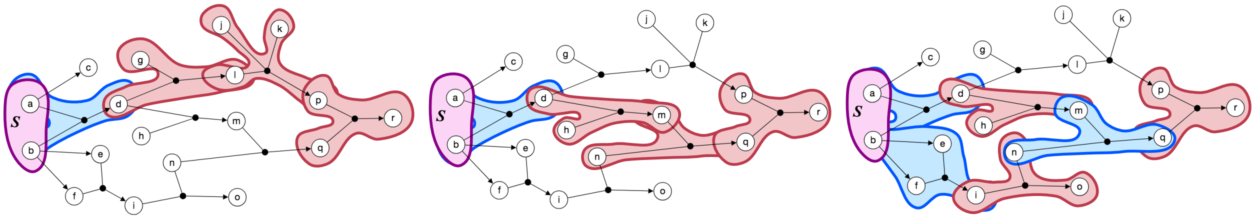

**Fig S10.** Examples of connectivity from $S = \{a, b\}$ to $r$ with a $B$-relaxation distance of three. Blue hyperedges denote traversals that are consistent with $B$-connectivity; red hyperedges denote traversals where one, but not all, nodes in the tail are connected; only hyperedges that are involved in the connectivity from $S$ to $r$ are highlighted for simplicity. Note that while $B$-relaxation distance is three, there are different sets of hyperedges that achieve this $B$-relaxation distance.

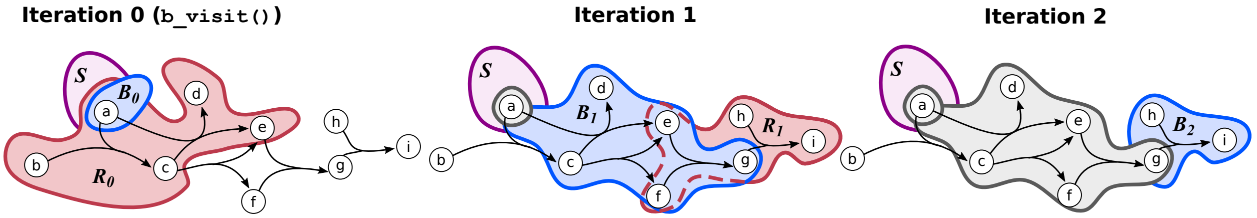

**Fig S11.** The restrictive set $R_k$ may include hyperedges that have been traversed in a previous iteration's `b_visit()` call. In iteration 1, the restrictive set $R_1$ is established by considering the $B$-connectivity from the heads of the two hyperedges in $R_0$. The hyperedge $\{\{e, f\}, \{g\}\}$ is restrictive with respect to the heads of one hyperedge in $R_0$ but traversable with respect to the heads of the other hyperedge. Thus, $\{\{e, f\}, \{g\}\}$ is included in $R_1$ but also added to the `seen` dictionary, saving redundant computation in Algorithm 2.
